# Identification of CRF66_BF, a new HIV-1 circulating recombinant form of South American origin

**DOI:** 10.1101/2021.09.14.460288

**Authors:** Joan Bacqué, Elena Delgado, Sonia Benito, María Moreno-Lorenzo, Vanessa Montero, Horacio Gil, Mónica Sánchez, María Carmen Nieto-Toboso, Josefa Muñoz, Miren Z. Zubero-Sulibarria, Estíbaliz Ugalde, Elena García-Bodas, Javier E. Cañada, Jorge del Romero, Carmen Rodríguez, Iciar Rodríguez-Avial, Luis Elorduy-Otazua, José J. Portu, Juan García-Costa, Antonio Ocampo, Jorge J. Cabrera, Michael M. Thomson

## Abstract

Circulating recombinant forms (CRFs) are important components of the HIV-1 pandemic. Among 108 reported in the literature, 17 are BF1 intersubtype recombinant, most of which are of South American origin. Among these, all 5 identified in the Southern Cone and neighboring countries, except Brazil, derive from a common recombinant ancestor related to CRF12_BF, which circulates widely in Argentina, as deduced from coincident breakpoints and clustering in phylogenetic trees. In a HIV-1 molecular epidemiological study in Spain, we identified a phylogenetic cluster of 20 samples from 3 separate regions which were of F1 subsubtype, related to the Brazilian strain, in protease-reverse transcriptase (Pr-RT) and of subtype B in integrase. Remarkably, 14 individuals from this cluster (designated BF9) were Paraguayans and only 4 were native Spaniards. HIV-1 transmission was predominantly heterosexual, except for a subcluster of 6 individuals, 5 of which were men who have sex with men. Ten additional database sequences, from Argentina (n=4), Spain (n=3), Paraguay (n=1), Brazil (n=1), and Italy (n=1), branched within the BF9 cluster. To determine whether it represents a new CRF, near full-length genome (NFLG) sequences were obtained for 6 viruses from 3 Spanish regions. Bootscan analyses showed a coincident BF1 recombinant structure, with 5 breakpoints, located in p17^gag^, integrase, gp120, gp41-*rev* overlap, and *nef*, which was identical to that of two BF1 recombinant viruses from Paraguay previously sequenced in NFLGs. Interestingly, none of the breakpoints coincided with those of CRF12_BF. In a maximum likelihood phylogenetic tree, all 8 NFLG sequences grouped in a strongly supported clade segregating from previously identified CRFs and from the CRF12_BF ‘family’ clade. These results allow us to identify a new HIV-1 CRF, designated CRF66_BF. Through a Bayesian coalescent analysis, the most recent common ancestor of CRF66_BF was estimated around 1984 in South America, either in Paraguay or Argentina. Among Pr-RT sequences obtained by us from HIV-1-infected Paraguayans living in Spain, 14 (20.9%) of 67 were of CRF66_BF, suggesting that CRF66_BF may be one of the major HIV-1 genetic forms circulating in Paraguay. CRF66_BF is the first reported non-Brazilian South American HIV-1 CRF_BF unrelated to CRF12_BF.

## 1. Introduction

One of the most distinctive features of HIV-1 evolution is its high recombinogenic potential, possibly the greatest among human pathogens, which is reflected in the high frequency of unique recombinant forms (URFs), each generated in a dually- or multiply-infected individual, found wherever different genetic forms circulate in the same population (Nájera et al., 2002). Some of the HIV-1 recombinant forms have spread beyond a group of epidemiologically-linked individuals, in which case they are designated circulating recombinant forms (CRFs) (Robertson et al., 2000). Currently, 108 CRFs have been reported in the literature and their number is increasing incessantly, due to both the generation of new CRFs and the identification of old previously undocumented CRFs. The proportion of CRFs in the HIV-1 pandemic has increased over time, representing around 17% infections in 2010-2015 (Hemelaar et al., 2020). Among CRFs, the most numerous are those derived from subtype B and subusbtype F1, 17 of which have been identified, most of them originated in South America, derived from the F1 variant circulating in Brazil (Louwagie et al., 1994). The first CRF_BF identified in South America was CRF12_BF, which circulates widely in Argentina and Uruguay, where URFs related to CRF12_BF are frequently found (Thomson et al., 2000; Thomson et al., 2002; Carr et al., 2001). Subsequently, 4 CRF_BFs related to CRF12_BF, as evidenced by shared breakpoints and phylogenetic clustering, were identified in the Southern Cone of South America or neighboring countries, CRF17_BF (Aulicino et al., 2012), CRF38_BF (Ruchansky et al., 2009), CRF44_BF (Delgado et al., 2010), and CRF89_BF (Delgado et al., 2021), the last three having clear country associations, with Uruguay, Chile, and Bolivia and Peru, respectively. Due to their common ancestry, these 5 CRFs and related URFs have been proposed to constitute a ‘family’ of recombinant viruses (Thomson and Nájera, 2005; Zhang et al., 2010; Delgado et al., 2021). By contrast, all CRF_BFs identified in Brazil are unrelated to CRF12_BF (De Sá Filho et al., 2006; Guimaraes et al., 2008; Sanabani et al., 2010; Pessoa et al., 2014; Reis et al., 2017; Reis et al., 2019). Here we report the identification of a new CRF_BF, found mainly in Paraguayan immigrants in Spain and also identified in Paraguay and Argentina. Interestingly, unlike all South American CRF_BFs identified to date outside of Brazil, it has no relationship with CRF12_BF.

## 2. Materials and methods

### 2.1. Samples

Plasma samples from HIV-1-infected individuals were collected in 14 Spanish regions for antiretroviral drug resistance tests and for a molecular epidemiological study. The study was approved by the Committee of Research Ethics of Instituto de Salud Carlos III, Majadahonda, Madrid, Spain. The study did not require written informed consent by the study participants, as it used samples and data collected as part of routine clinical practice and patients’ data were anonymized without retaining data allowing individual identification.

### 2.2. RNA extraction, RT-PCR amplification, and sequencing

An ~1.4 kb *pol* fragment in protease-reverse transcriptase (Pr-RT) was amplified from plasma RNA by RT-PCR followed by nested PCR as described previously (Delgado et al., 2015) and sequenced with the Sanger method using a capillary automated sequencer. Some samples were also subject to amplification and sequencing of integrase. Near full-length genome (NFLG) sequences were obtained for selected samples by RT-PCR/nested PCR amplification from plasma RNA in four overlapping segments and sequenced by the Sanger method, as described (Delgado et al., 2002; Sierra et al., 2005; Cañada et al., 2021). Newly derived sequences are deposited in GenBank under accessions MK298150, OK011530-OK011552.

### 2.3. Phylogenetic sequence analyses

Sequences were aligned with MAFFT v7 (Katoh and Standley, 2013). Initial phylogenetic trees with all Pr-RT sequences obtained by us were constructed via approximate maximum likelihood with FastTree v2.1.10 (Price et al., 2010), using the general time reversible evolutionary model with CAT approximation to account for among-site rate heterogeneity, with assessment of node support with Shimodaira-Hasegawa (SH)-like local support values (Guindon et al., 2010). Subsequent maximum likelihood (ML) trees with sequences of interest were constructed with W-IQ-Tree (Trifinopoulos et al., 2016), using the best-fit substitution model selected by ModelFinder program (Kalyaanamoorthy et al., 2017), with assessment of node support with the ultrafast bootstrap approximation approach (Hoang et al., 2018). Trees were visualized with MEGA v7.0 (Kumar et al., 2016).

Mosaic structures were analyzed by bootscanning (Salminen et al., 1995) with SimPlot v1.3.5 (Lole et al., 1999). In these analyses, trees were constructed using the neighbor-joining method with the Kimura 2-parameter model and a window width of 400 nucleotides. Recombinant segments identified with SimPlot were further phylogenetically analyzed via ML with W-IQ-Tree. Intersubtype breakpoint locations were also determined with jpHMM (Schultz et al., 2009).

### 2.4. Temporal and geographical estimations

The time and the location of the most recent common ancestor (MRCA) of the identified CRF was estimated using Pr-RT sequences with the Bayesian Markov chain Monte Carlo (MCMC) coalescent method implemented in BEAST v1.10.4 (Suchard et al., 2018). Prior to the BEAST analysis, the existence of temporal signal in the dataset was assessed with TempEst v1.5.3 (Rambaut et al., 2016), which determines the correlation of genetic divergence among sequences (measured as root-to-tip distance) with time. The BEAST analysis was performed using the SRD06 codon-based evolutionary model (with two codon position partitions: 1st+2nd and 3rd) (Shapiro et al., 2006). We also specified an uncorrelated lognormal relaxed clock and a Bayesian SkyGrid coalescent tree prior (Gill et al., 2013). The MCMCs were run for 50 million generations. We performed runs in duplicate, combining the posterior tree files with LogCombiner v1.10.4. Mixing and convergence were checked with Tracer v1.6, ensuring that effective sample size values of all parameters were >200. The posterior distribution of trees was summarized in a maximum clade credibility (MCC) tree with TreeAnnotator v1.10.4, after removal a 10% burn-in. MCC trees were visualized with FigTree v1.4.2 (Rambaut, http://tree.bio.ed.ac.uk/software/figtree/). Parameter uncertainty was summarized in 95% highest posterior density (HPD) intervals.

## 3. Results

### 3.1. Identification of a BF recombinant cluster and epidemiological associations

In a molecular epidemiology study on HIV-1 in Spain we identified a cluster of 20 viruses of F1 subsubtype in Pr-RT, that in integrase, sequenced in 4 samples, were of subtype B, which was designated BF9. Inclusion in the phylogenetic analyses of Pr-RT sequences of all viruses in the Los Alamos HIV Sequence Database (Los Alamos National Laboratory, 2021) classified as being of F1 subsubtype or BF1 recombinant identified 10 additional viruses belonging to BF9, from Argentina (n=4), Spain (n=3), Paraguay (n=1), Brazil (n=1), and Italy (n=1) (Fig. 1). Pr-RT sequences of the BF9 cluster were most closely related to F1 viruses of the Brazilian variant. Epidemiological data of the 20 samples of the BF9 cluster from Spain processed by us are shown in Table 1. Remarkably, 14 individuals were from Paraguay and all 3 remaining database sequences from samples collected in Spain were from Latin Americans, one each from Paraguay, Argentina, and an unspecified country. Transmission was predominantly heterosexual, but 7 were men who have sex with men (MSM), the sequences of 5 of whom branched in a subcluster (Fig. 1).

**Figure 1.**
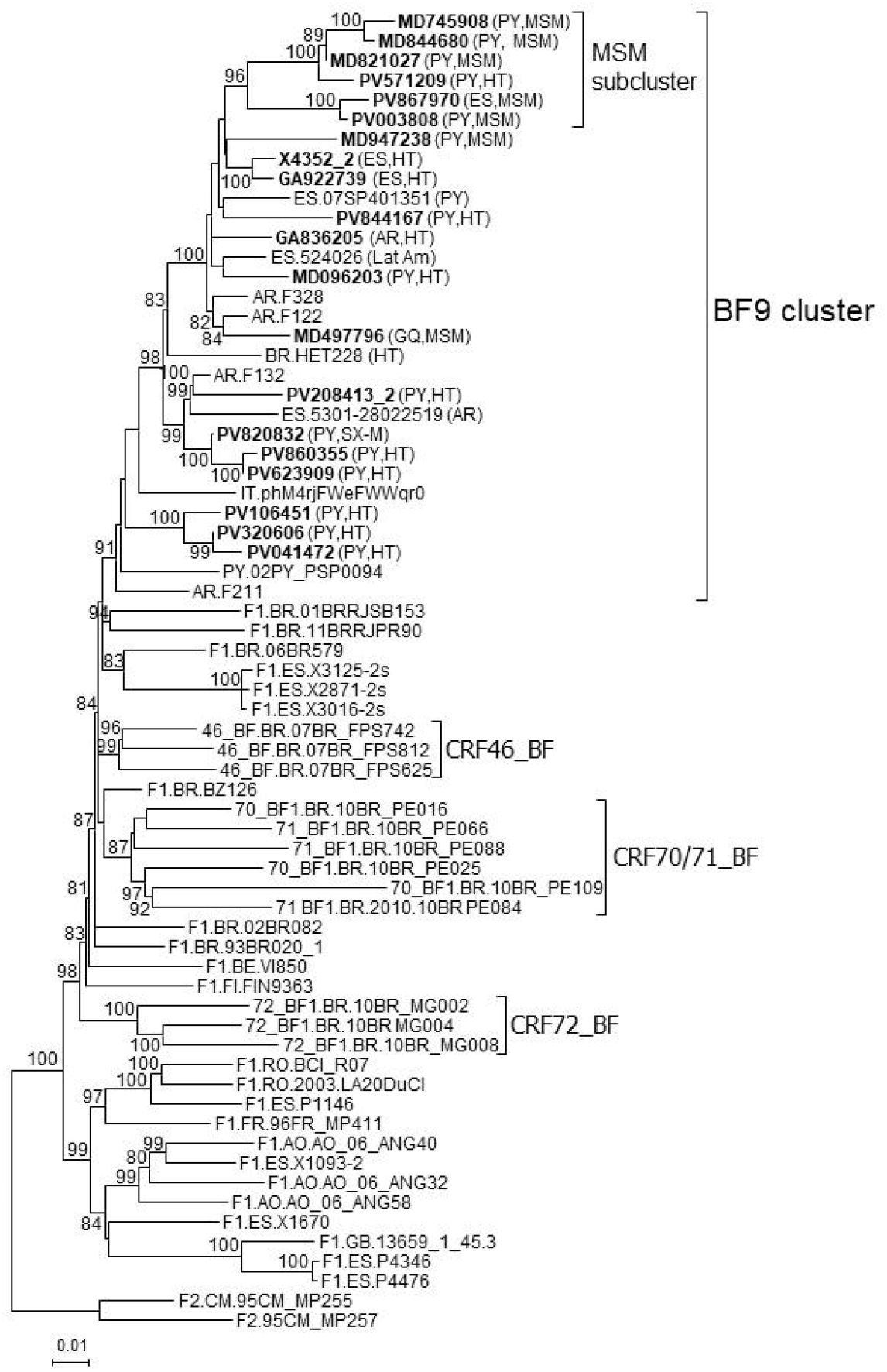
Maximum likelihood tree of Pr-RT sequences of BF9 cluster. Names of sequences obtained by us, all collected in Spain, are in bold type. In database sequences, the country of sample collection is indicated before the virus name with the 2-letter ISO country code. After the names of viruses of the BF9 cluster, the 2-letter ISO code of country of origin of the patient and/or the transmission route, when known, are shown in parentheses. F1 sequences from different countries are included in the analysis, together with two F2 sequences used as outgroups. Only ultrafast bootstrap values ≥80% are shown. PY: Paraguay; AR: Argentina; ES: Spain; BR: Brazil; IT: Italy; GQ: Equatorial Guinea; MSM: man who has sex with men; HT: heterosexual; SX-M: male with unspecified sexual acquisition of HIV-1.

**Table 1.** Epidemiological data of patients and GenBank accessions of sequences.

| **SAMPLE ID** | **City of sample collection** | **Region of sample collection** | **Year of sample collection** | **Year of HIV diagnosis** | **Gender** | **Transmission route\*** | **Country of origin** | **GenBank accessions** |
| --- | --- | --- | --- | --- | --- | --- | --- | --- |
| **X4352_2** | Vigo | Galicia | 2017 | 2017 | M | HT | Spain | MK298150 (NFLG) |
| **GA836205** | Vigo | Galicia | 2020 | 2020 | M | HT | Argentina | OK011531 (Pr-RT)<br>OK011530 (integrase) |
| **GA922739** | Ourense | Galicia | 2018 | 2017 | M | HT | Spain | OK011532 (NFLG) |
| **MD096203** | Madrid | Madrid | 2017 | 2011 | F | HT | Paraguay | OK011534 (Pr-RT)<br>OK011533 (integrase) |
| **MD497796** | Madrid | Madrid | 2017 | 2017 | M | MSM | Equatorial Guinea | OK011534 (NFLG) |
| **MD745908** | Madrid | Madrid | 2019 | 2019 | M | MSM | Spain | OK011536 (Pr-RT) |
| **MD821027** | Madrid | Madrid | 2018 | 2018 | M | MSM | Paraguay | OK011537 (Pr-RT) |
| **MD844680** | Madrid | Madrid | 2020 | 2020 | M | MSM | Paraguay | OK011538 (Pr-RT) |
| **MD947238** | Madrid | Madrid | 2018 | 2016 | M | MSM | Paraguay | OK011539 (Pr-RT) |
| **PV003808** | Bilbao | Basque Country | 2020 | 2020 | M | MSM | Paraguay | OK011541 (Pr-RT)<br>OK011540 (integrase) |
| **PV041472** | Bilbao | Basque Country | 2014 | 2014 | F | HT | Paraguay | OK011542 (Pr-RT) |
| **PV106451** | Bilbao | Basque Country | 2010 | 2010 | F | HT | Paraguay | OK011543 (NFLG) |
| **PV208413_2** | Bilbao | Basque Country | 2009 | 2009 | M | HT | Paraguay | OK011544 (Pr-RT) |
| **PV320606** | Bilbao | Basque Country | 2014 | 2014 | M | HT | Paraguay | OK011545 (Pr-RT) |
| **PV571209** | Bilbao | Basque Country | 2013 | 2013 | M | HT | Paraguay | OK011546 (Pr-RT) |
| **PV623909** | Bilbao | Basque Country | 2011 | 2011 | F | HT | Paraguay | OK011547 (NFLG) |
| **PV820832** | Bilbao | Basque Country | 2008 | 2008 | M | Sexual | Paraguay | OK011548 (Pr-RT) |
| **PV844167** | Vitoria | Basque Country | 2016 | 2016 | M | HT | Paraguay | OK011549 (NFLG) |
| **PV860355** | Bilbao | Basque Country | 2011 | 2011 | M | HT | Paraguay | OK011550 (Pr-RT) |
| **PV867970** | Bilbao | Basque Country | 2020 | 2020 | M | MSM | Spain | OK011552 (Pr-RT)<br>OK011551 (integrase) |

\*HT: heterosexual; MSM: man who has sex with men.

### 3.2. Analyses of NFLG sequences and identification of a new CRF

To determine whether viruses from the BF9 cluster represent a new CRF, we obtained NFLG sequences from 6 samples from 3 Spanish regions and analyzed their mosaic structures by bootscanning. Two additional NFLG sequences of BF recombinant viruses from databases were also analyzed by bootscanning, both from Paraguay: 02PY_PSP0094, that branched in the BF9 cluster in Pr-RT, and 02PY_PSP0093, that showed high similarity to NFLGs of the BF9 cluster in BLAST searches of the Los Alamos database. All 8 analyzed genomes showed coincident mosaic structures, with 5 breakpoints, located in p17^gag^, integrase, gp120, gp41-*rev* overlap, and *nef* (Fig. 2). Breakpoints were more precisely located using the midpoint of B-F1 transitions, according to the positions where 75% consensuses of subtype B and the F1 Brazilian strain genomes differ, in HXB2 positions 950, 4327, 6486, 8498, and 9161. Breakpoint locations were also determined with jpHMM (Supplementary Table), which also found 5 breakpoints for each virus in intervals overlapping those of the other analyzed viruses and the 75% consensus B-F1 transition intervals in all cases except the breakpoint interval in *nef* of MD497796, that did not overlap the consensus B-F1 transition interval, and that in p17^gag^ of PV106451, that was not detected by jpHMM. ML phylogenetic trees constructed with each interbreakpoint fragment confirmed the subtype assignation determined with bootscanning (Fig. 3).

**Figure 2.**
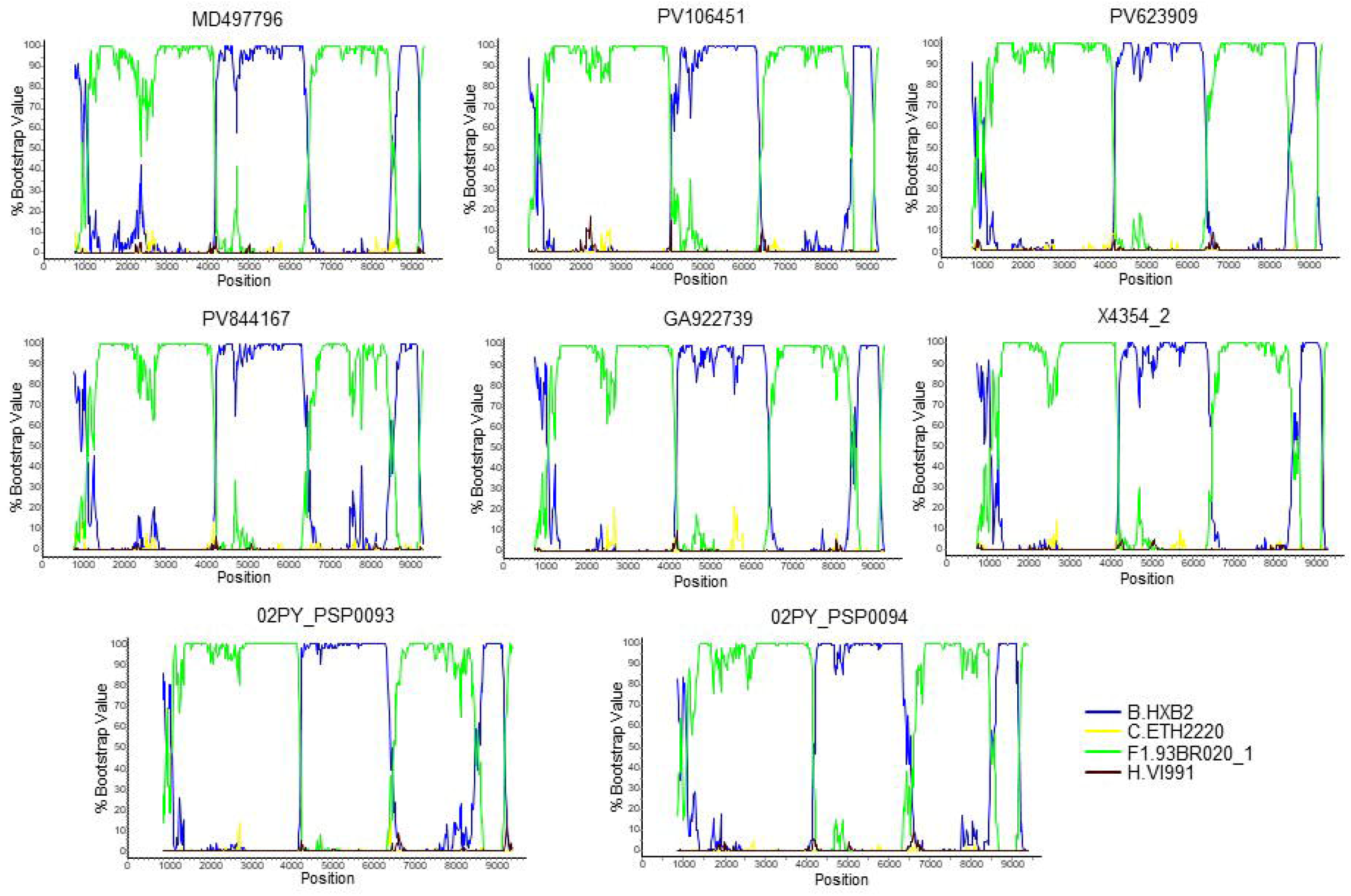
Bootscan analyses of 6 NFLG sequences of viruses of the BF9 cluster obtained by us and of two BF1 database NFLG sequences from Paraguay, 02PY_PSP0093 and 02PY_PSP0094. The horizontal axis represents the position in the HXB2 genome of the midpoint of a 400 nt window moving in 20 nt increments and the vertical axis represents bootstrap values supporting clustering with subtype reference sequences.

**Figure 3.**
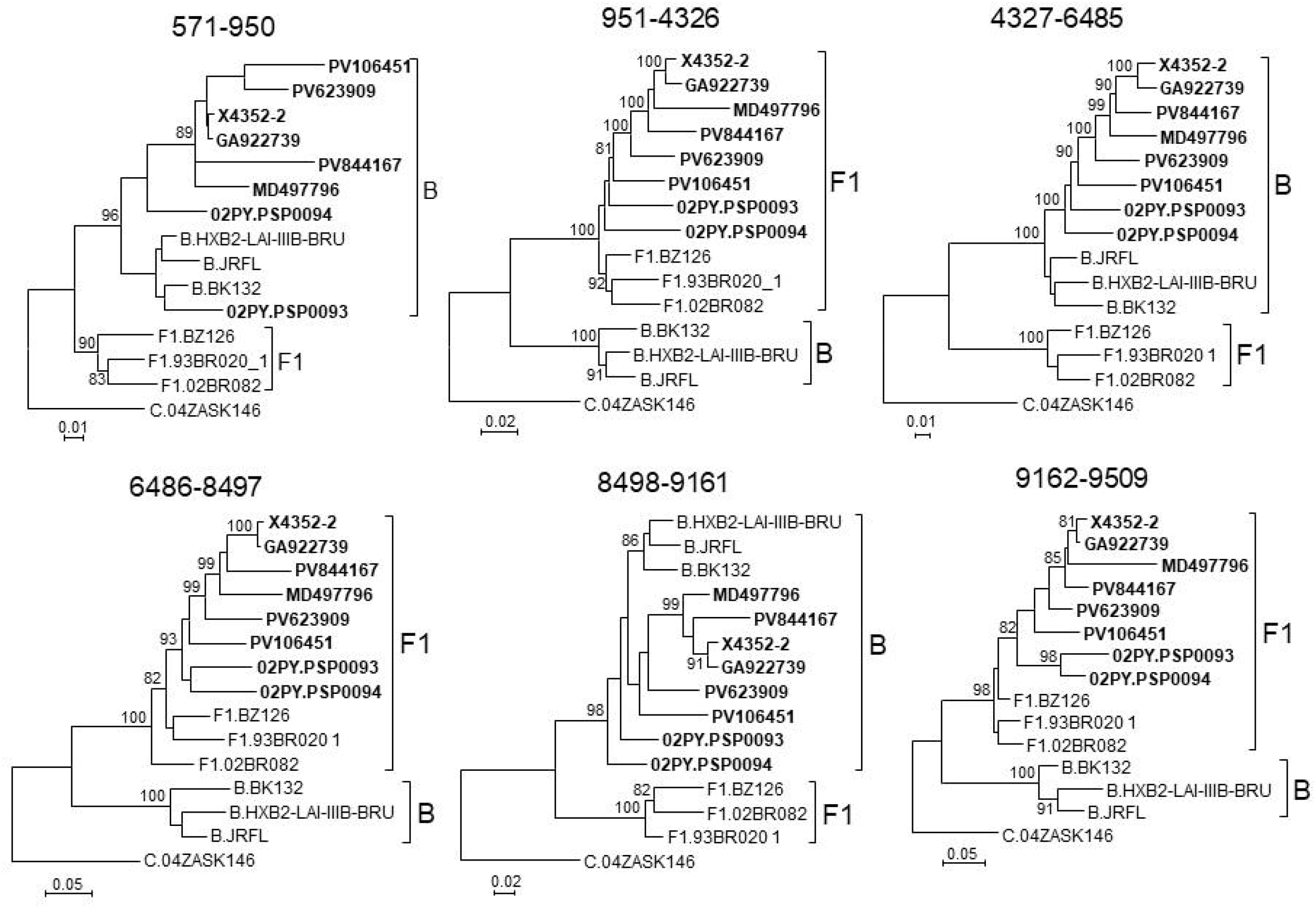
Phylogenetic trees of interbreakpoint genome segments of the BF recombinant viruses analyzed by bootscanning. HXB2 positions delimiting the analyzed segments are indicated on top of the trees. Sequence names of BF viruses are in bold type. Names of subtype references are preceded by the corresponding subtype name. Only ultrafast bootstrap values ≥80% are shown.

In an ML tree constructed with the 7 NFLG genomes of the BF9 cluster and 02PY_PSP0093, all 8 genomes grouped in a strongly supported clade segregating away from all other CRF_BFs and of the clade formed by the 5 CRF_BFs of the CRF12_BF family (Fig. 4). It should be pointed out that 02PY_PSP0093 did not branch in the BF9 cluster in the tree of Pr-RT, which suggests that the Pr-RT segment of this virus could derive from secondary recombination with an F1 strain different from the parental F1 strain of all other BF9 viruses.

**Figure 4.**
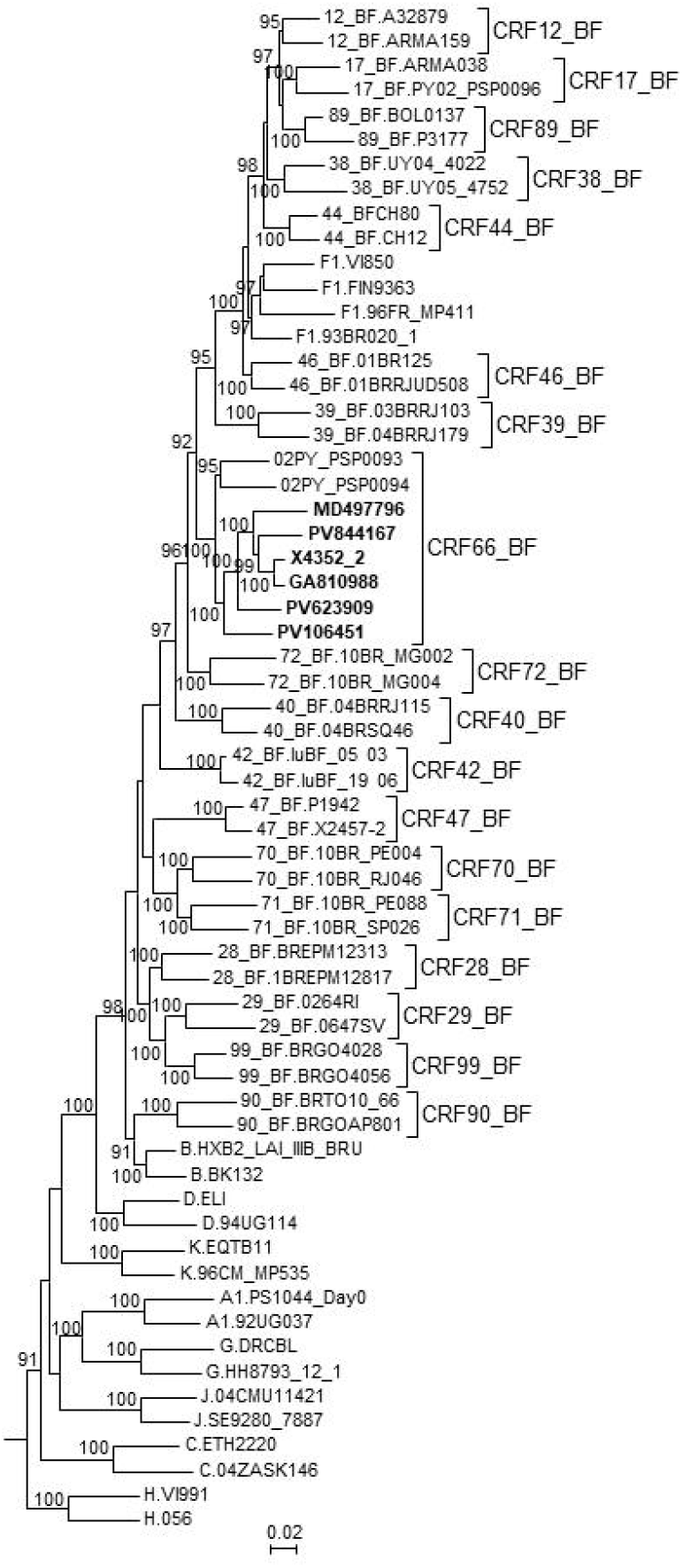
Maximum likelihood tree of NFLG sequences of viruses of the BF9 cluster and PY02_PSP0094. References of all published CRF_BFs and of HIV-1 subtypes are also included in the analysis. The tree is rooted with SIVcpz virus MB66. Names of sequences obtained by us are in bold type. In reference sequences, the subtype or CRF is indicated before the virus name. Only ultrafast bootstrap values ≥90% are shown.

These results, therefore, allow to define a new CRF, which was designated CRF66_BF, whose mosaic structure is shown in Fig. 5.

**Figure 5.**
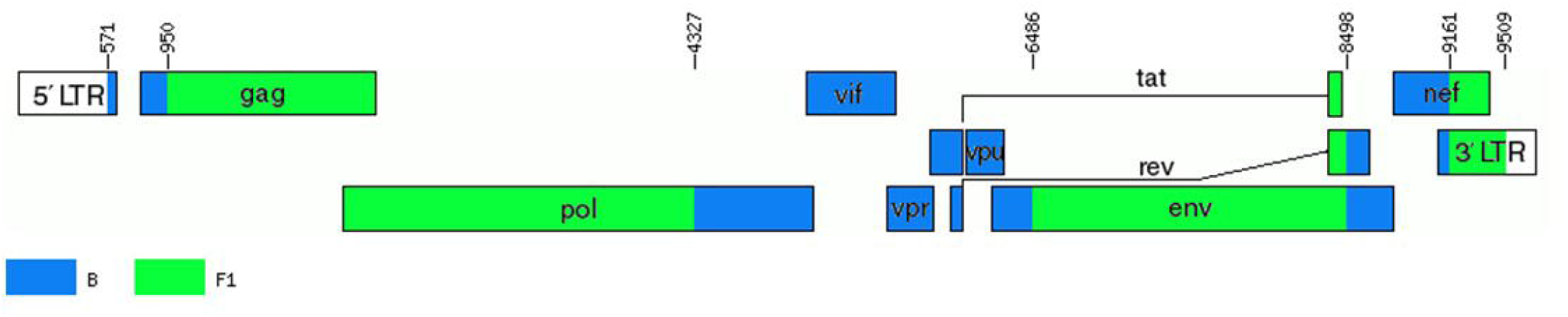
Mosaic structure of CRF66_BF. Breakpoint positions are numbered as in the HXB2 genome. The drawing was made using the Recombinant HIV-1 Drawing Tool https://www.hiv.lanl.gov/content/sequence/DRAW_CRF/recom_mapper.html

### 3.3. Prevalence of CRF66_BF among HIV-1-infected Paraguayans residing in Spain

Among 67 HIV-1-infected Paraguayans residing in Spain studied by us, CRF66_BF was the most common non-subtype B genetic form, representing 20.9% (14 of 67) of total infections, 48.3% (14 of 29) of non-subtype B infections, and 60.1% (14 of 23) of F1/BF1 infections.

### 3.4. Temporal and geographical estimations of CRF66_BF origin

To estimate the time and place of origin of CRF66_BF, Pr-RT sequences where analyzed with a Bayesian coalescent method with BEAST 1.10.4. Prior to this analysis, TempEst analysis determined that there was an adequate temporal signal in the dataset (r^2^ = 0.5871). In the BEAST analysis, for the sequences corresponding to South American individuals residing in Spain, the assigned location trait was their country of origin, rather than their place of residence. This was done because most individuals harboring CRF66_BF identified in Spain were of South American origin (mostly from Paraguay) and because we found no definitive evidence of the local circulation of CRF66_BF in Spain, as reflected in clusters mainly comprising Spanish individuals. Therefore, we assumed that South Americans harboring CRF66_BF viruses had acquired HIV-1 in their countries of origin. In this analysis, the time of the MRCA of CRF66_BF was estimated around 1984 (95% HPD, 1971-1996), and its most probable location was in Paraguay (PP=0.77), with Argentina in second place (PP=0.20) (Fig. 6). Considering the possibility that local subclusters each found in one city could represent local transmissions, we performed a second analysis in which we assigned the country of location of the most recent diagnoses of such clusters to Spain, irrespective of the countries of origin of the individuals. In this analysis, Paraguay was also estimated as the most probable location of the MRCA of CRF66_BF, although with a lower support (PP=0.55), and the support for Argentina increased to a PP=0.42 (Supplementary Figure 1).

**Figure 6.**
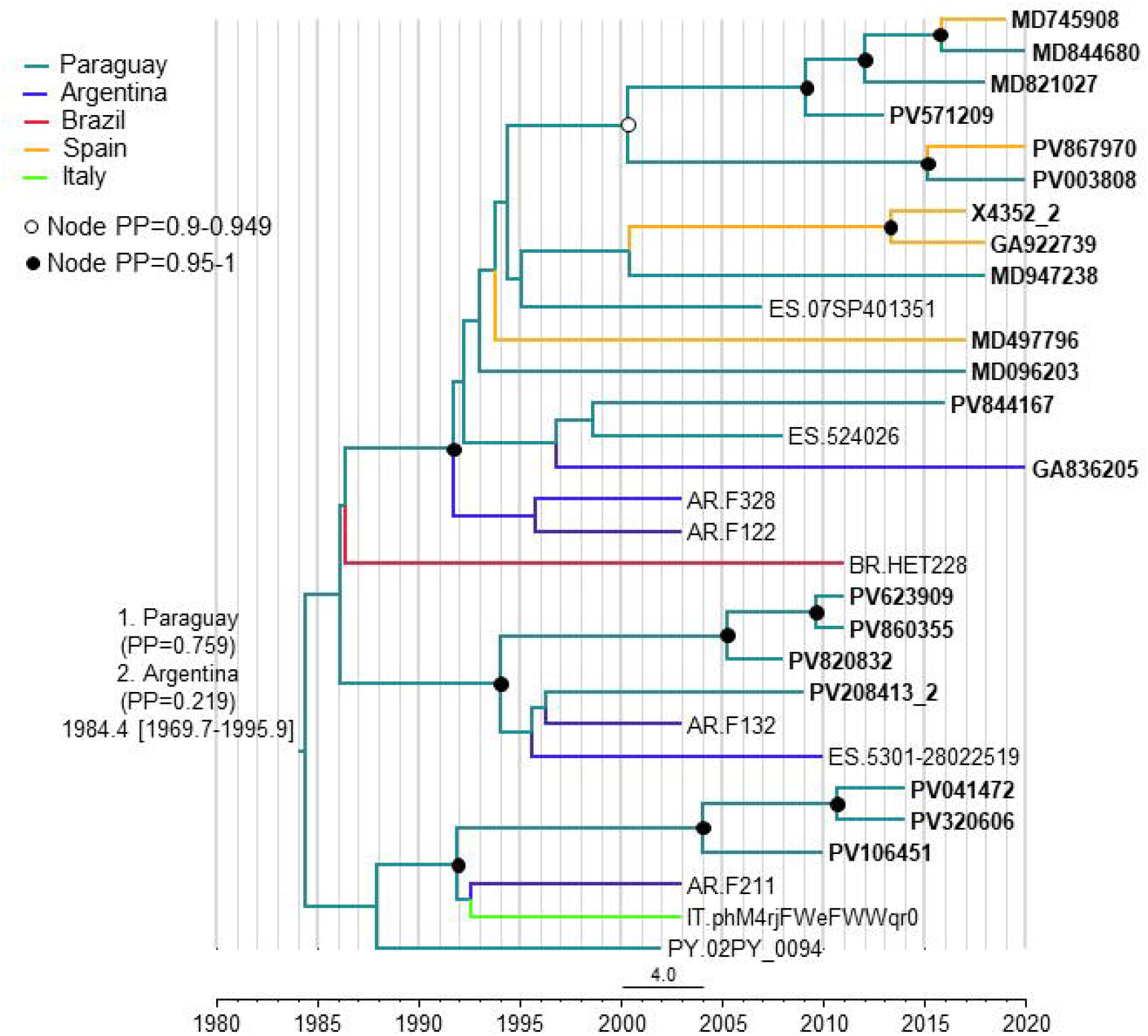
Maximum clade credibility tree of CRF66_BF Pr-RT sequences. Branch colors indicate, for terminal branches, country of sample collection or, for South American individuals residing in Spain, of origin of the individual, which was used as location trait (see Methods), and for internal branches, the most probable location country of the subtending node, according to the legend on the upper left. For database sequence 524026, from a sample collected in Spain, location was assigned to Paraguay as the most probably country of origin, although the only available information in the GenBank entry is that the individual was from Latin America, because 15 (88.2%) of 17 Latin Americans with CRF66_BF sampled in Spain were from Paraguay. Nodes supported by PP≥0.95 and PP 0.9-0.949 are indicated with filled and unfilled circles, respectively. The two most probable countries at the root of the tree are indicated, together with the PPs supporting each location and the time of the MRCA (mean value, with 95% HPD interval in brackets).

## 4. Discussion

The results of this study allow to define a new HIV-1 CRF, designated CRF66_BF, which is the 18^th^ reported CRF derived from subtypes B and F. Samples harboring CRF66_BF were collected in 5 countries, in South America (Argentina, Paraguay, and Brazil) and Western Europe (Spain and Italy), with a majority collected in Spain. However, of samples collected in Spain, a great majority were from Paraguayan individuals. Bayesian coalescent analyses (performed with the assumption that South American individuals living in Spain harboring CRF66_BF viruses had acquired them in their countries of origin), pointed to a most probable origin of CRF66_BF in Paraguay (PP=0.77), with Argentina being the second most probable location (PP=0.20). When the analysis was performed assigning the most recently diagnosed samples of clusters found in a single Spanish city to Spain as the location trait, irrespective of the country of origin of the individual, the PPs for a MRCA in Paraguay or Argentina were not very different (0.55 vs. 0.42, respectively). Therefore, the results point to a South American origin of CRF66_BF, either in Paraguay or Argentina, without a definitive support for either country. However, given the great predominance of Paraguayans among CRF66_BF-infected individuals living in Spain, we cannot rule out that the same could happen in Argentina, where Paraguayans represent the largest immigrant national group (Instituto Nacional de Estadísticas y Censos, República Argentina, 2021). If this was the case, and information on country of origin of the sampled individuals living in Argentina was included in the analyses, it is possible that the support for a root of the CRF66_BF tree in Paraguay would increase.

Among HIV-1-infected Paraguayans residing in Spain studied by us, there was relatively high prevalence (21%) of CRF66_BF infections, which suggests that CRF66_BF could be one of the major HIV-1 genetic forms circulating in Paraguay. A better knowledge of the current prevalence of CRF66_BF in Paraguay would require sequencing a representative sample of recent HIV-1 diagnoses in the country. However, HIV-1 sequences from only 24 patients sampled in Paraguay are available at the Los Alamos HIV Sequence database, and the most recent molecular epidemiological study published to date involves the analysis of sequences from 55 samples collected 18 to 19 years ago (Aguayo et al., 2008), which are not available in public databases.

Notably, CRF66_BF, unlike all other non-Brazilian CRF_BFs identified to date in South America (CRF12_BF, CRF17_BF, CRF38_BF, CRF44_BF, and CRF89_BF, all circulating in the Southern Cone or neighboring countries), is unrelated to CRF12_BF, as deduced from the lack of breakpoint coincidence and of phylogenetic clustering with CRF12_BF. This implies that CRF66_BF originated independently from viruses of the CRF12_BF family, with a presumable ancestry in Brazil, where B and F1 viruses are circulating (Louwagie et al., 1994).

CRF66_BF is the 5th CRF of South American ancestry originally identified in Western Europe [after CRF42_BF (Struck et al., 2015), CRF47_BF (Fernández-García et al., 2010), CRF60_BC (Simonetti et al., 2014), and CRF89_BF (Delgado et al., 2021)], which, together with the reported propagation of HIV-1 variants of South America origin among the European population (Delgado et al., 2015; de Oliveira et al., 2010; Collaço Verás et al., 2012; Thomson et al., 2012; Vinken et al., 2019; Lai et al., 2014; Fabeni et al., 2015; Fabeni et al., 2020; Carvalho et al., 2015), points to an increasing relationship between the HIV-1 epidemics in both continents. This reflects migratory fluxes, most notably in Spain, where around 2.5 million South Americans live, representing nearly 40% of the migrant population (Instituto Nacional de Estadística, 2021a), and immigration from South America has increased greatly in recent years (Instituto Nacional de Estadística, 2021b) (Supplementary Figure 2). Considering the large and increasing South American immigrant population in Europe and the scarcity of HIV-1 sequences from most South American countries available in public databases, studies on HIV-1 genetic diversity and molecular epidemiology among South American immigrants living in Europe could provide novel insights into the HIV-1 epidemics in their countries of origin, as well as on the diffusion of South American HIV-1 variants in Europe.

It is interesting to note that although transmission of CRF66_BF is predominantly via heterosexual contact, most individuals in a cluster are MSM (Fig. 1), which suggests diffusion from a heterosexual to a MSM network. HIV-1 propagation between heterosexual and MSM networks has also been reported for CRF89_BF (Delgado et al., 2021) and for a large CRF02_AG cluster in Spain (Delgado et al., 2019), although in the latter case the direction of propagation was from MSM to heterosexuals.

One of the essential tasks of Biology is naming and classifying organisms. In this work, we have accomplished this task by identifying a new HIV-1 circulating recombinant form, derived from subtypes B and F1, named CRF66_BF. CRF66_BF most likely originated in South America, either in Paraguay or Argentina, and, unlike all non-Brazilian South American CRFs identified to date, is unrelated to CRF12_BF. The identification and genetic characterization of HIV-1 variants is the first and necessary step for molecular epidemiological studies examining their geographic dissemination, growth dynamics, and epidemiological associations, as well as for analyzing their biological properties, such as pathogenic and transmissibility potentials, response to antiretroviral therapies, and susceptibility to immune responses inducible by vaccines.

## Acknowledgments

We thank José Antonio Taboada, from Consellería de Sanidade, Xunta de Galicia, and Daniel Zulaika, from Osakidetza-Servicio Vasco de Salud, for their support of this study, and the personnel at the Genomic Unit, Instituto de Salud Carlos III, for technical assistance in sequencing.

## Author contributions

MT and ED conceived the study and supervised the experimental work. JB, MT, and ED processed sequences and performed phylogenetic analyses. MT performed phylodynamic analyses. HG performed data curation. JB, SB, MM-L, VM, MS, EG-B, and JEC performed experimental work. MCN-T, JM, MZZ-S, EU, JdR, CR, IR-A, LE-O, JJP, JG-C, AO, and JJC obtained samples and epidemiological data from patients. MT, ED, and HG wrote the manuscript. All authors read and approved the manuscript.

