## Supplementary figures and images for "Identification of CRF66_BF, a new HIV-1 circulating recombinant form of South American origin"

### Supplementary Figure 1

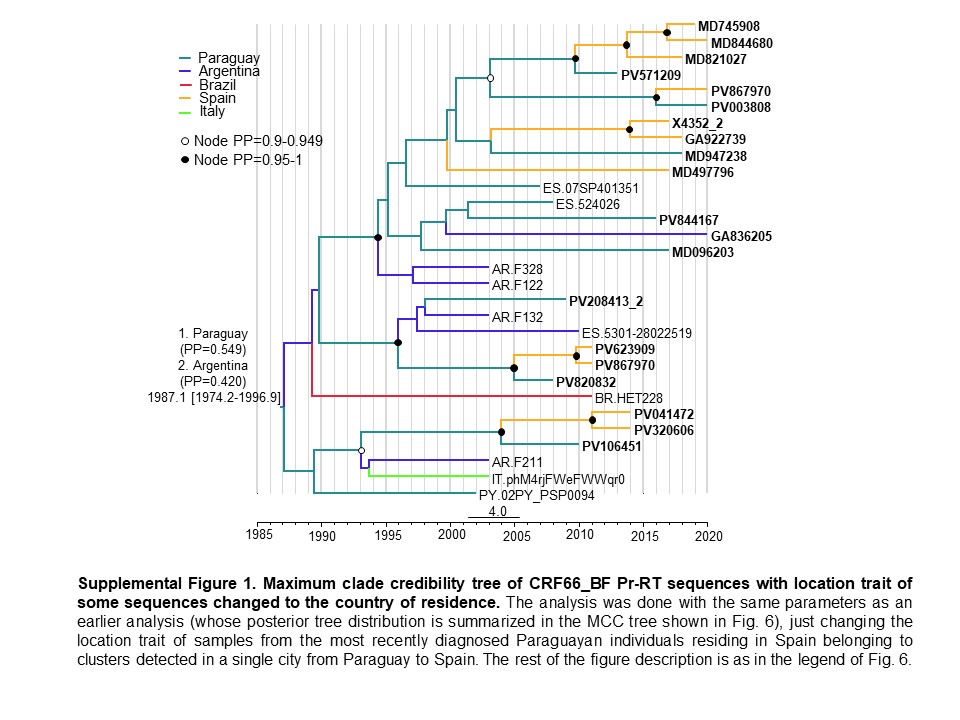

### Supplementary Figure 2

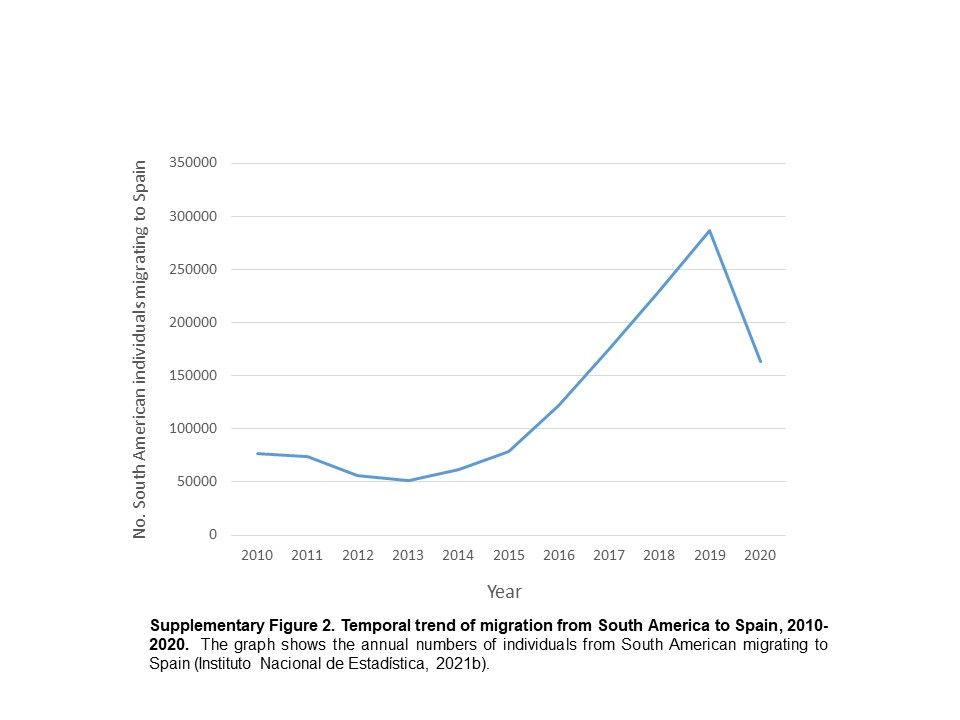
