## Supplementary Table for "Identification of CRF66_BF, a new HIV-1 circulating recombinant form of South American origin"

| **Supplementary Table. Breakpoints (HXB2 positions) detected by jpHMM in the analyzed BF recombinant genome sequences.** | | | | | |
| --- | --- | --- | --- | --- | --- |
| **Sample ID** | **Interclade B-F1 consensus transition midpoint and interval** | | | | |
|  | 950  (940-960) | 4327  (4311-4342) | 6486  (6481-6490) | 8498  (8487-8508) | 9161  (9107-9215) |
|  | **jpHMM breakpoints** | | | | |
| MD497796 | 950±10 | 4312±30 | 6532±51 | 8474±28 | 9309±19† |
| PV106451 | N.D.* | 4353±56 | 6509±28 | 8495±9 | 9176±67 |
| PV623909 | 950±12 | 4312±30 | 6563±82 | 8474±30 | 9178±110 |
| PV844167 | 1027±71 | 4301±41 | 6565±85 | 8481±8 | 9244±76 |
| GA922739 | 1025±71 | 4308±34 | 6532±51 | 8482±7 | 9260±61 |
| X4352_2 | 979±28 | 4308±34 | 6531±50 | 8482±7 | 9255±65 |
| 02PY_PSP0093 | 951±10 | 4310±33 | 6511±26 | 8490±14 | 9311±101 |
| 02PY_PSP0094 | 938±21 | 4318±46 | 6492±11 | 8483±18 | 9161±51 |
| *N.D.: breakpoint not detected. †The breakpoint interval determined by jpHMM does not overlap the consensus interclade B-F1 interval. | | | | | |
